## Supplementary Material for "Abundance and diversity of resistomes differ between healthy human oral cavities and gut"

### Supplementary Figures

a

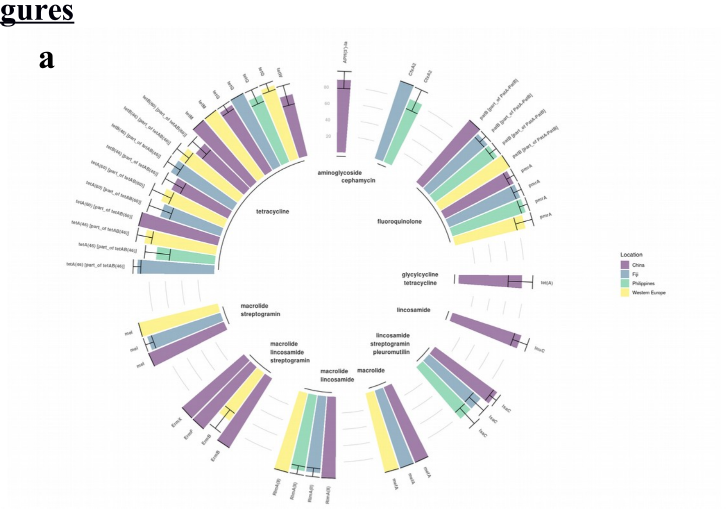

b

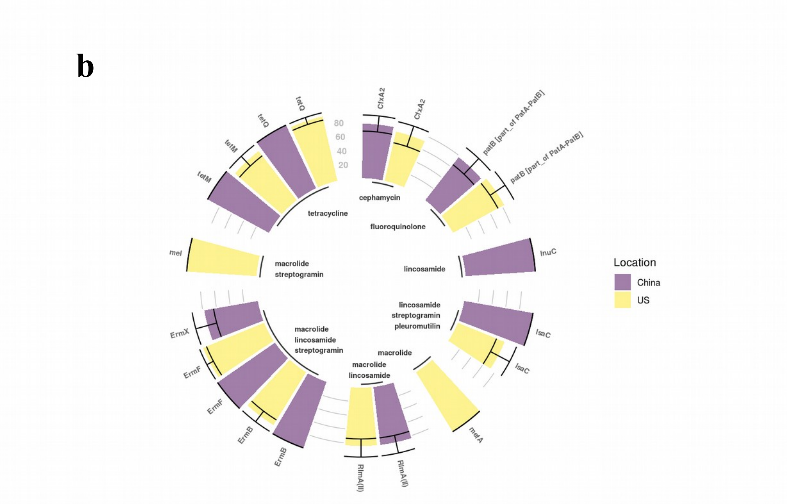


c

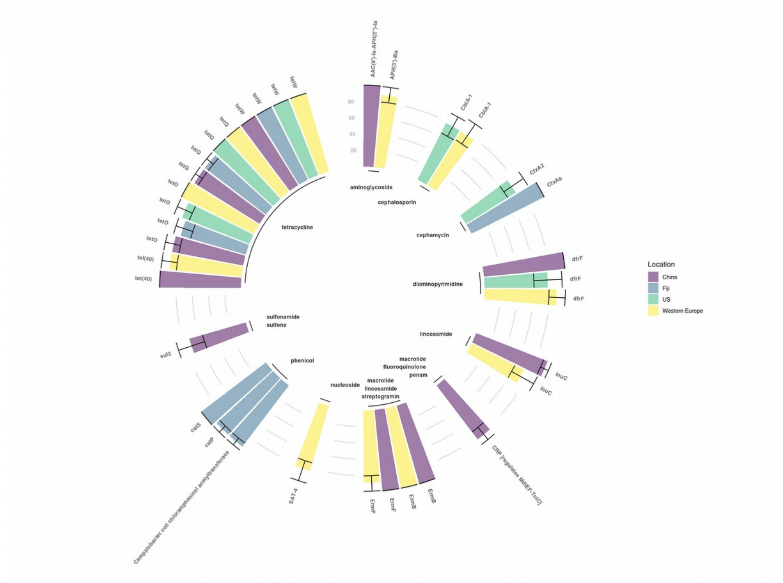

**Supplementary Fig. 1** ARGs, grouped by ARG class, that are found >= 70% of samples for **a** saliva from China (n = 18), Fiji (n = 18), the Philippines (n = 18) and Western Europe (n = 18), **b** dental plaque from China (n = 18) and the US (n = 18), and **c** stool from China (n = 18), Fiji (n = 18), the US (n = 18) and Western Europe (n = 18). Error bars are 95 % confidence intervals (CIs) that were evaluated from percentages extracted from bootstrapping samples 100 times.

**Supplementary Fig. 2** Dendrogram of only longitudinal US samples where abundance (log10[RPKM+1]) is clustered by hierarchical clustering (complete method on Euclidean distance matrix) and labelled by sample type (buccal mucosa: n = 150, dorsum of tongue: n = 173, dental plaque: n = 178, stool n = 148). These samples were collected within two years with individuals having had no antimicrobial treatment in that time.

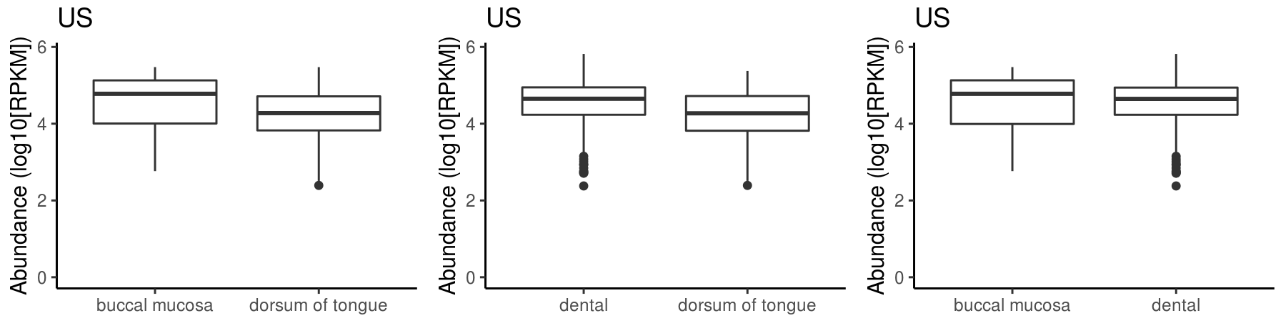


**Supplementary Fig. 3** Absolute abundance in log10 of reads per kilobase of read per million (RPKM) of ARGs for paired samples between US buccal mucosa (n = 86) and dorsum of tongue (n = 86), US dental plaque (n = 89) and dorsum of tongue (n = 89), and US buccal mucosa (n = 86) and dental plaque (n = 86)

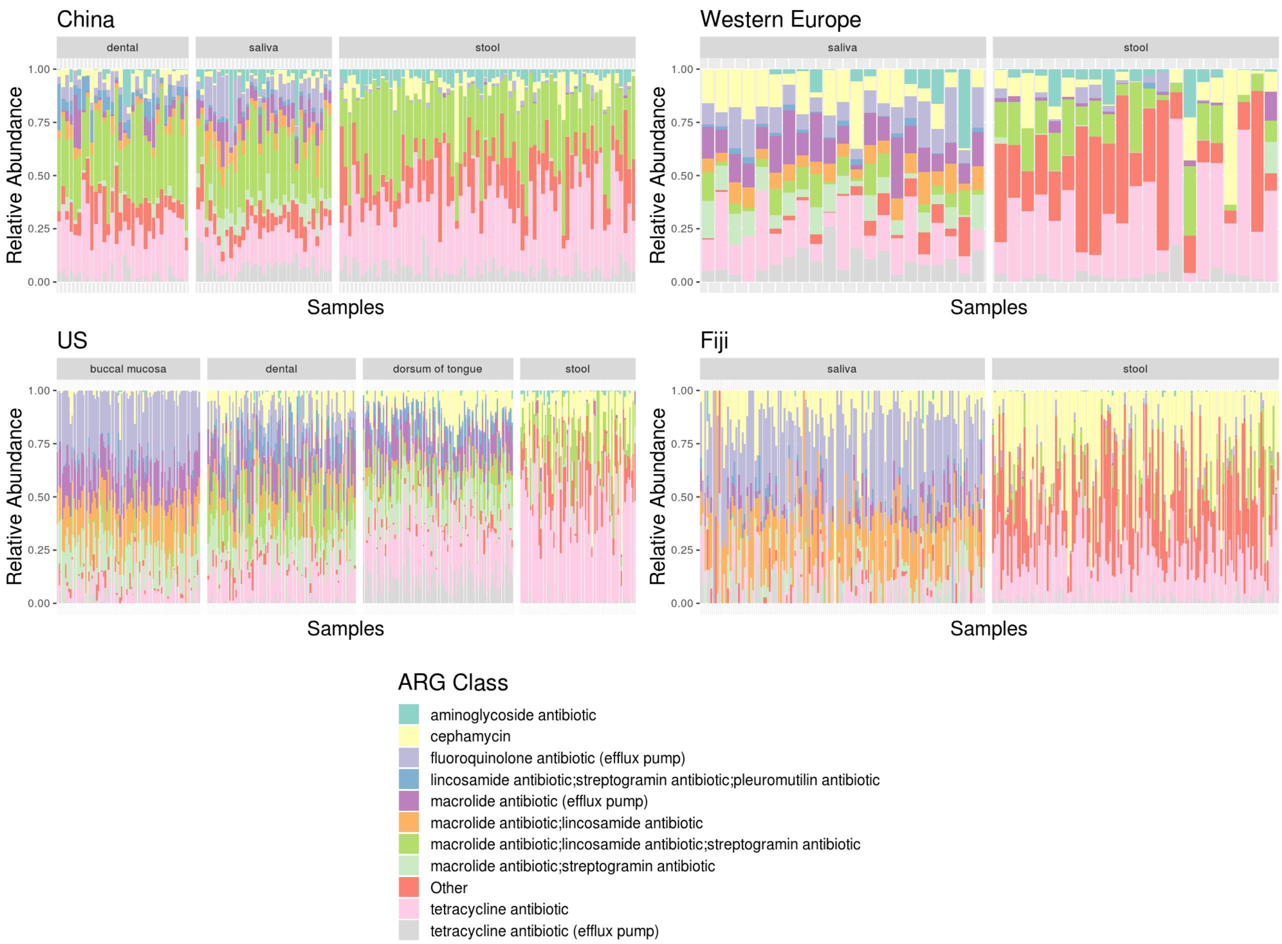


**Supplementary Fig. 4** Relative abundance of reads labelled by the top ten most abundant ARG classes across all geographical locations or "Other" classes for each sample from China saliva (n = 33), dental plaque (n = 32) and stool (n = 72), Western Europe saliva (n = 21) an stool (n = 21), US buccal mucosa (n = 87), dental plaque (n = 90), dorsum of tongue (n = 91) and stool (n = 70), and Fiji saliva (n = 136) and stool (n = 137)

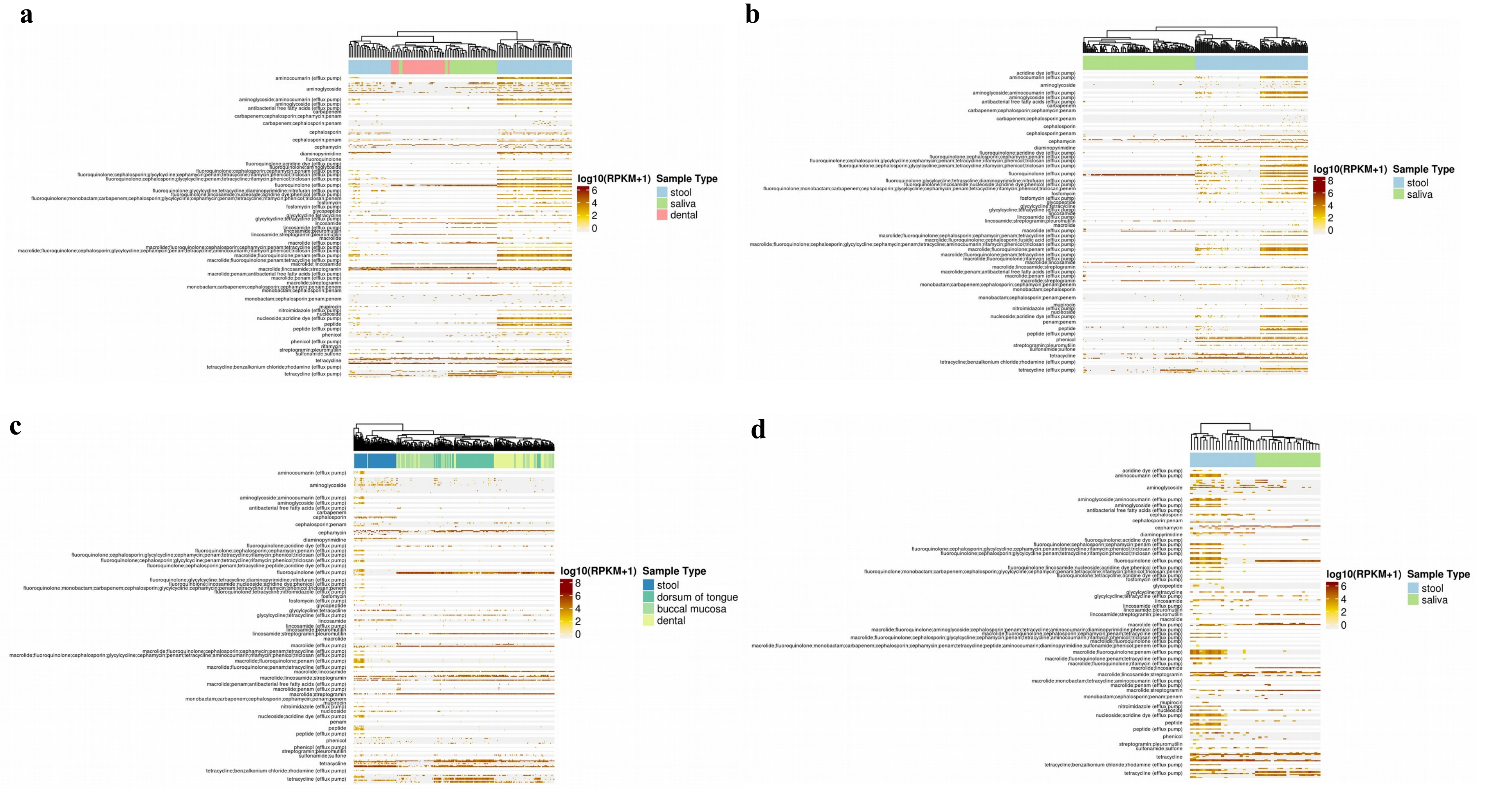


**Supplementary Fig. 5** Heatmaps of abundance (log10[RPKM+1]) clustered by hierarchical clustering column-wise by sample (complete method on euclidean distance matrix) and separated row-wise by ARG class for **a** China (saliva: n = 33, dental plaque: n = 32, stool: n = 72); **b** US including longitudinal samples (buccal mucosa: n = 164, dorsum of tongue: n = 188, dental plaque: n = 191, stool n = 156); **c** Fiji (saliva: n = 136, stool: n = 137); and **d** Western Europe (saliva: n = 21, stool: n = 21)

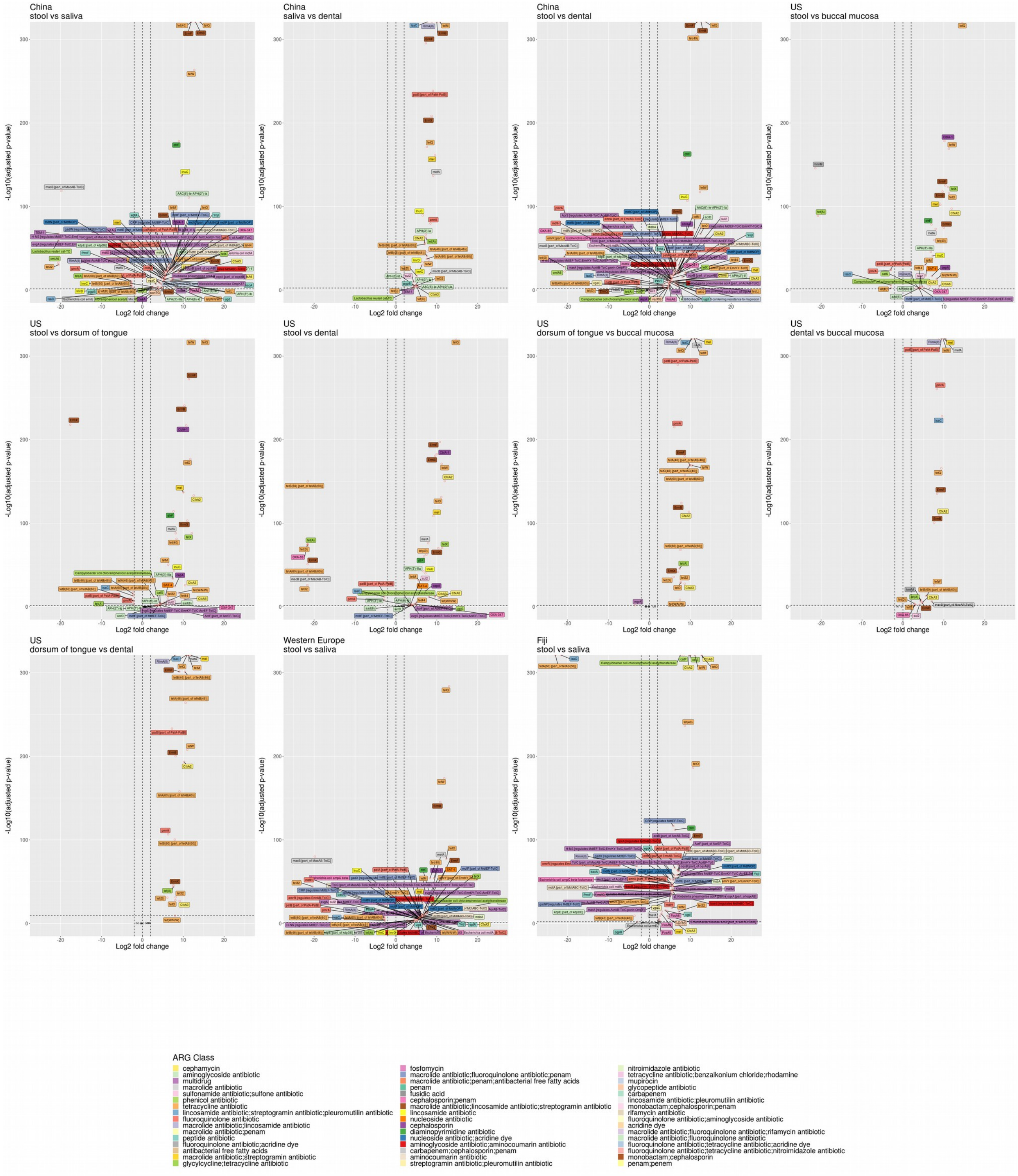


**Supplementary Fig. 6** Volcano plots showing differential analysis, using DESeq2 package in R, between paired samples of adjusted p-value < 0.05 between: China stool (n = 31) and saliva (n = 31); China dental plaque (n = 31) and saliva (n = 31); China stool (n = 30) and dental plaque (n = 30); US stool (n = 64) and buccal mucosa (n = 64); US stool (n = 69) and dorsum of tongue (n = 69); US stool (n = 68) and dental plaque (n = 68); US dorsum of tongue (n = 86) and buccal mucosa (n = 86); US buccal mucosa (n = 86) and dental plaque (n = 86); US dorsum of tongue (n = 89) and dental plaque (n = 89); Western Europe saliva (n = 21) and stool (n = 21); Fiji saliva (n = 132) and stool (n = 132).

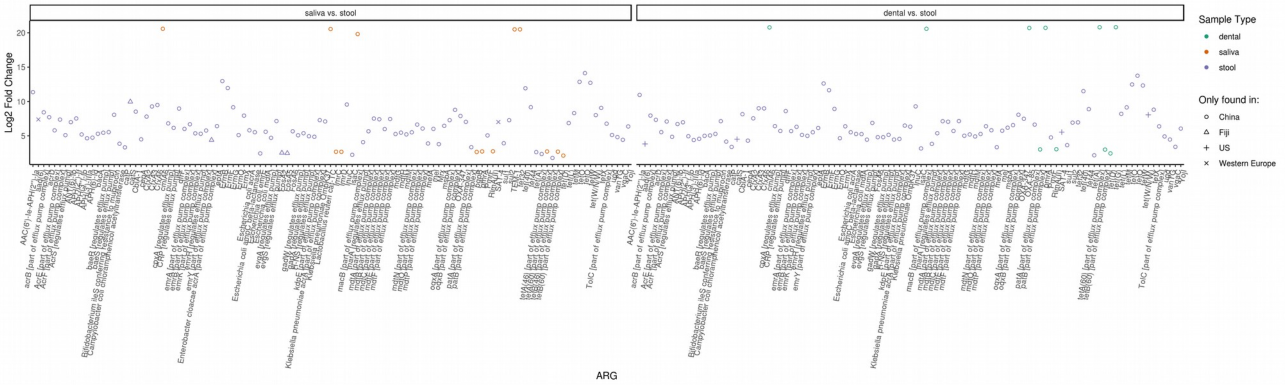


**Supplementary Fig. 7** Log2 fold change of ARGs exclusively found in one geographical location between paired China stool (n = 31) and saliva (n = 31), Fiji saliva (n = 137) and stool (n = 137), Western Europe saliva (n = 21) and stool (n = 21), China stool (n = 30) and dental plaque (n = 30), and US stool (n = 68) and dental plaque (n = 68) samples (adjusted p-value < 0.05).

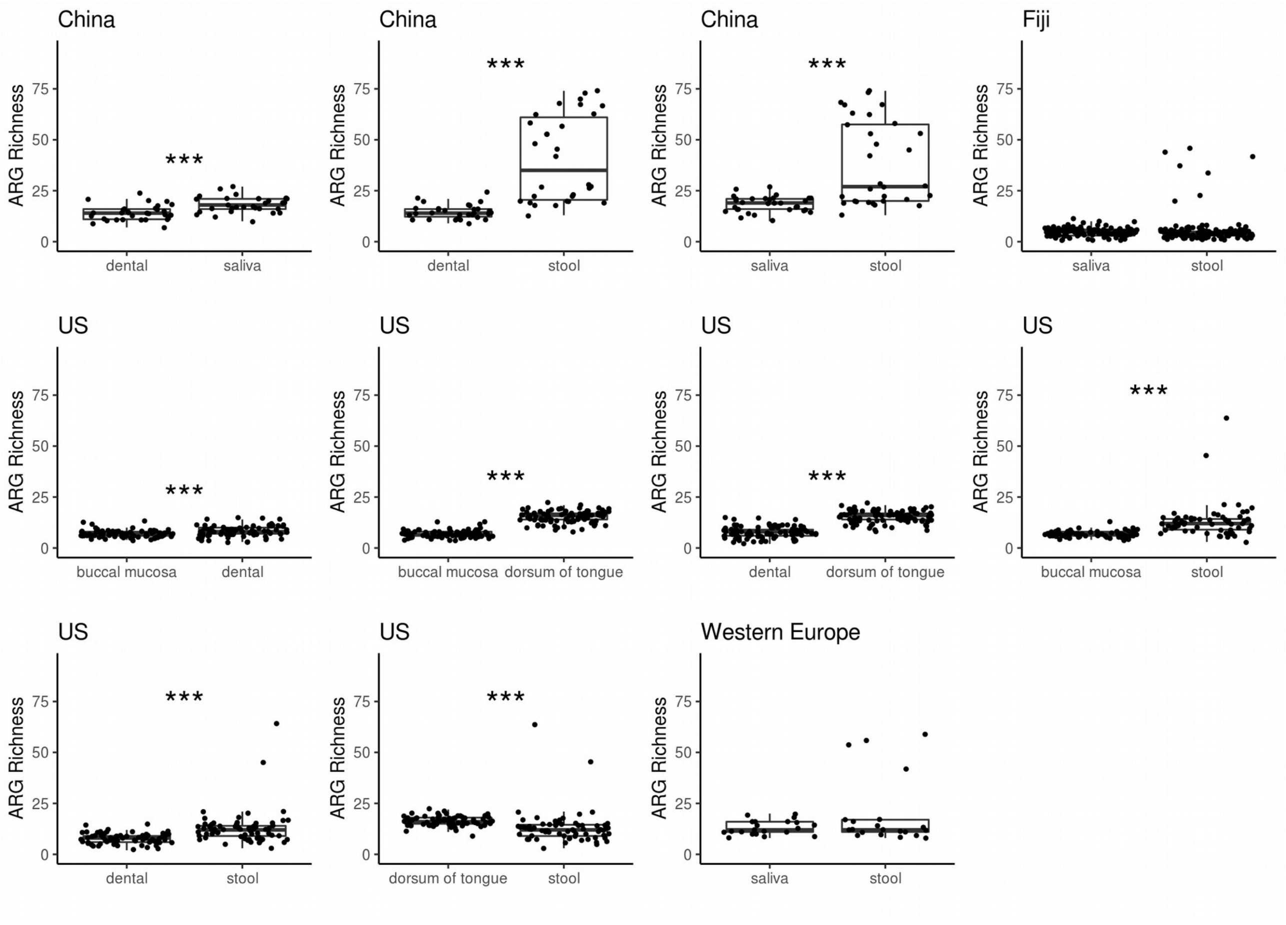


**Supplementary Fig. 8** ARG richness for paired samples, with ARGs that are part of or regulate an efflux pump complex removed, between China dental plaque (n = 31) and saliva (n = 31), China stool (n = 30) and dental plaque (n = 30), China stool (n = 31) and saliva (n = 31), Fiji saliva (n = 128) and stool (n = 128), US buccal mucosa (n = 78) and dental plaque (n = 78), US buccal mucosa (n = 86) and dorsum of tongue (n = 86), US dental plaque (n = 89) and dorsum of tongue (n = 89), US buccal mucosa (n = 64) and stool (n = 64), US dental plaque (n = 68) and stool (n = 68), US dorsum of tongue (n = 67) and stool (n = 67), and Western Europe saliva (n = 21) and stool (n = 21) with Mann-Whitney, paired, two-sided t-test (p-value < 0.05 as *, < 0.01 as **, < 0.005 as ***)

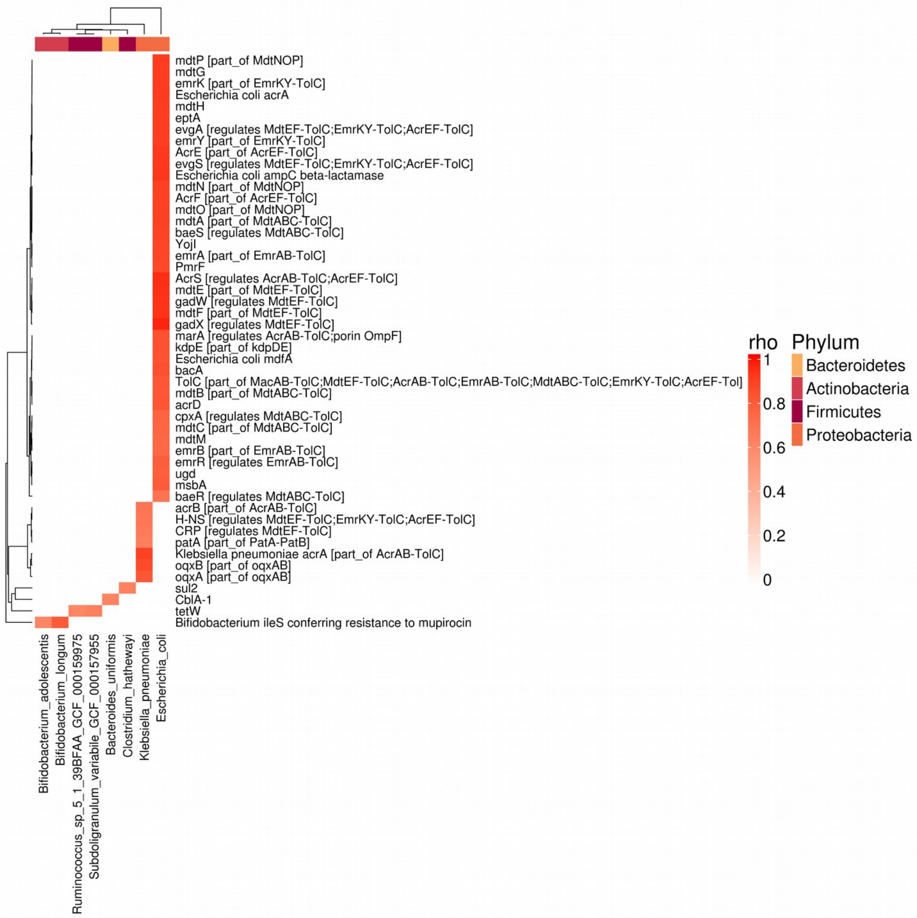


**Supplementary Fig. 9** Spearman's correlation of ARG and species abundance from China stool samples (n = 31) represent as a heatmap of *rho* between 0 and 1, only where adjusted p-value < 0.05. Rows and columns are clustered by hierarchical clustering of Euclidean distance. Columns are coloured by phylum. P-values are adjusted by Benjamini-Hochberg multiple test correction.
